# Identifying the gene responsible for NPQ reversal in *Phaeodactylum tricornutum*

**DOI:** 10.1101/2024.03.27.587055

**Authors:** Maxwell A. Ware, Andrew J. Paton, Yu Bai, Tessema Kassaw, Martin Lohr, Graham Peers

## Abstract

Algae such as diatoms and haptophytes have distinct photosynthetic pigments from plants, including a novel set of carotenoids. This includes a primary xanthophyll cycle comprised of diadinoxanthin and its de-epoxidation product diatoxanthin that enables the switch between light harvesting and non-photochemical quenching (NPQ)-mediated dissipation of light energy. The enzyme responsible for the reversal of this cycle was previously unknown. Here, we identified zeaxanthin epoxidase 3 (ZEP3) from *Phaeodactylum tricornutum* as the candidate diatoxanthin epoxidase. Knocking out the ZEP3 gene caused a loss of rapidly reversible NPQ following saturating light exposure. This correlated with the maintenance of high concentrations of diatoxanthin during recovery in low light. Xanthophyll cycling and NPQ relaxation were restored via complementation of the wild type ZEP3 gene. The *zep3* knockout strains showed reduced photosynthetic rates at higher light fluxes and reduced specific growth rate in variable light regimes, likely due to the mutant strains becoming locked in a light energy dissipation state. We were able to toggle the level of NPQ capacity in a time and dose dependent manner by placing the ZEP3 gene under the control of an inducible promoter. Identification of this gene provides deeper understanding of the diversification of photosynthetic control in algae compared to plants and suggests a potential target to improve the productivity of industrial-scale cultures.

## Introduction

There are a diverse set of pigments used by plants and algae for photosynthesis and for photoprotection. Researchers focused on vascular plants are familiar with chlorophylls *a+b,* which are used for light energy capture (Kunugi et al., 2016). Conversely, the carotenoid pigments of the xanthophyll cycle (violaxanthin, antheraxanthin and zeaxanthin; VAZ) are known to play an important role in regulating the thermal dissipation of excess light energy through non-photochemical quenching (NPQ, (Jahns et al., 2009). The enzymes violaxanthin de-epoxidase (VDE) and zeaxanthin epoxidase (ZEP) contribute to toggling the light-harvesting antennae between a light-harvesting state and energy-dissipation state, respectively (Bugos and Yamamoto, 1996; Marin et al., 1996; Jahns et al., 2009). VDE removes the epoxide group from the ionone rings of violaxanthin to yield zeaxanthin which promotes NPQ. ZEP converts zeaxanthin back to violaxanthin thereby re-establishing efficient light utilization.

Many algae, examples of which include diatoms and diatoms, contain a different set of photosynthetic pigments from plants. The brown color of these algae is primarily due to the presence of chlorophylls *a+c* and the carotenoid fucoxanthin which participate in light harvesting (Büchel, 2020). Furthermore, the xanthophyll cycle of these algae utilize diadinoxanthin and diatoxanthin (Ddx:Dtx), which are absent from the green lineage of algae and plants (Takaichi, 2011; Bai et al., 2022). Under certain conditions, algae with the Ddx:Dtx cycle may also accumulate the pigments of the VAZ cycle, but probably not to a significant extent under most natural conditions (Lohr and Wilhelm, 1999).

NPQ is the major photoprotective mechanism in diatoms. Indeed, the rapidly inducible component of NPQ, qE, can dissipate 50% of absorbed photons as heat (Giovagnetti et al., 2014). The de-/epoxidation cycle works as follows: Ddx is the xanthophyll correlated with light harvesting under light limited conditions, and a high luminal pH. During high light induced low luminal pH, Ddx is de-epoxidated to Dtx. This promotes NPQ similar to zeaxanthin in the VAZ cycle (Arsalane et al., 1994). Gene silencing experiments suggested that the VDE gene is a key component for the conversion of Ddx to Dtx in diatoms (Lavaud et al., 2012). However, the reverse reaction has not been resolved.

Reverse genetics experiments facilitate the discovery of biosynthetic pathways associated with light harvesting and NPQ-related pigments in diatoms and related algae. It was recently discovered that a novel type of dioxygenase is responsible for the final biosynthesis step of chlorophyll *c* in diatoms and dinoflagellates. This enzyme is absent from plants and green algae (Jiang et al., 2023; Jinkerson et al., 2024). There has also been increased interest in uncovering the biosynthesis pathway of fucoxanthin. This pigment is an asymmetrical carotenoid containing unusual chemical features such as an allenic bond and a keto group in its polyene moiety. In diatoms and haptophytes, fucoxanthin biosynthesis is facilitated by a series of gene duplications and neofunctionalizations that occurred during/post-secondary endosymbiosis. For instance, a carotenoid isomerase-like protein CRTISO5 does not function as an isomerase. Instead, it hydrates a carbon-carbon triple bond, converting the intermediate phaneroxanthin to fucoxanthin (Cao et al., 2023). Moreover, paralogues of VDE and ZEP contribute to fucoxanthin biosynthesis. Violaxanthin De-Epoxidase-Like 1 (VDL1) catalyzes the tautomerization of violaxanthin to neoxanthin (Dautermann et al., 2020). VDL2 performs the tautomerization of diadinoxanthin to allenoxanthin (Bai et al., 2022). Meanwhile, the epoxidation of haptoxanthin to phaneroxanthin, the penultimate step of fucoxanthin biosynthesis, is catalyzed by ZEP1(Bai et al., 2022).

The genome of the model diatom *Phaeodactylum tricornutum* encodes two additional ZEP genes (ZEP2 and ZEP3, Coesel et al., 2008; Bai et al., 2022). We hypothesized that ZEP3 may be playing a role in the photoprotective Ddx:Dtx cycle as its expression has been found to be co-regulated with that of VDE (Nymark et al., 2013). Here we show that insertional knockout of this gene results in a reduced variable NPQ capacity due to the constitutive accumulation of diatoxanthin and reduced growth rates in variable light environments compared to WT strains.

## Results

### Knockout of PtZEP3

We hypothesized that the *Phaeodactylum* gene annotated as zeaxanthin epoxidase 3 (*Pt*ZEP3, Phatr3_J10970) was the diatoxanthin epoxidase. We targeted a knockout of this gene using CRISPR-Cas9 and homology directed repair-mediated insertion of a bleomycin resistance cassette. We employed two guide RNAs targeted to the 3’ end of exon 2 in the gene sequence (Fig S1). This approach resulted in the generation of several homozygous *zep3* knockouts that grew on antibiotic selection. ZEP3 gene disruptions were verified via PCR amplification of the ZEP3 gene and its increased size compared to the WT (Fig S1). Primers directed towards the *ble* antibiotic resistance gene and the 3’ end of the WT gene verified the disruption of ZEP3 in our KO and complemented strains, with no product in the WT. Complementation of a mutant with the native gene sequence restored the WT band, which has been observed previously for this methodology (Fig S1,Bai et al., 2022).

### ZEP3 is Necessary for Rapidly Reversible NPQ Capacity and Diatoxanthin Epoxidation

WT, *zep3* KO strains, and complemented strains were cultivated in a 12h light:12h dark sinusoidal light regime. We first tested their NPQ capacity within an hour after dawn. We reasoned that sampling shortly after dawn would allow for wild type *Phaeodactylum* to relax NPQ overnight. When we measured the NPQ capacity of WT *ex situ*, we found high NPQ capacity was rapidly induced with a saturating light intensity of 2000 μmol photons m^−2^ s^−1^. NPQ then relaxed within 30 minutes of low light (Fig 1A). In contrast, there was very little induced NPQ observed in both *Ptzep3* mutants and we observed little change throughout the low light relaxation phase (Fig 1A).

**Fig. 1.**
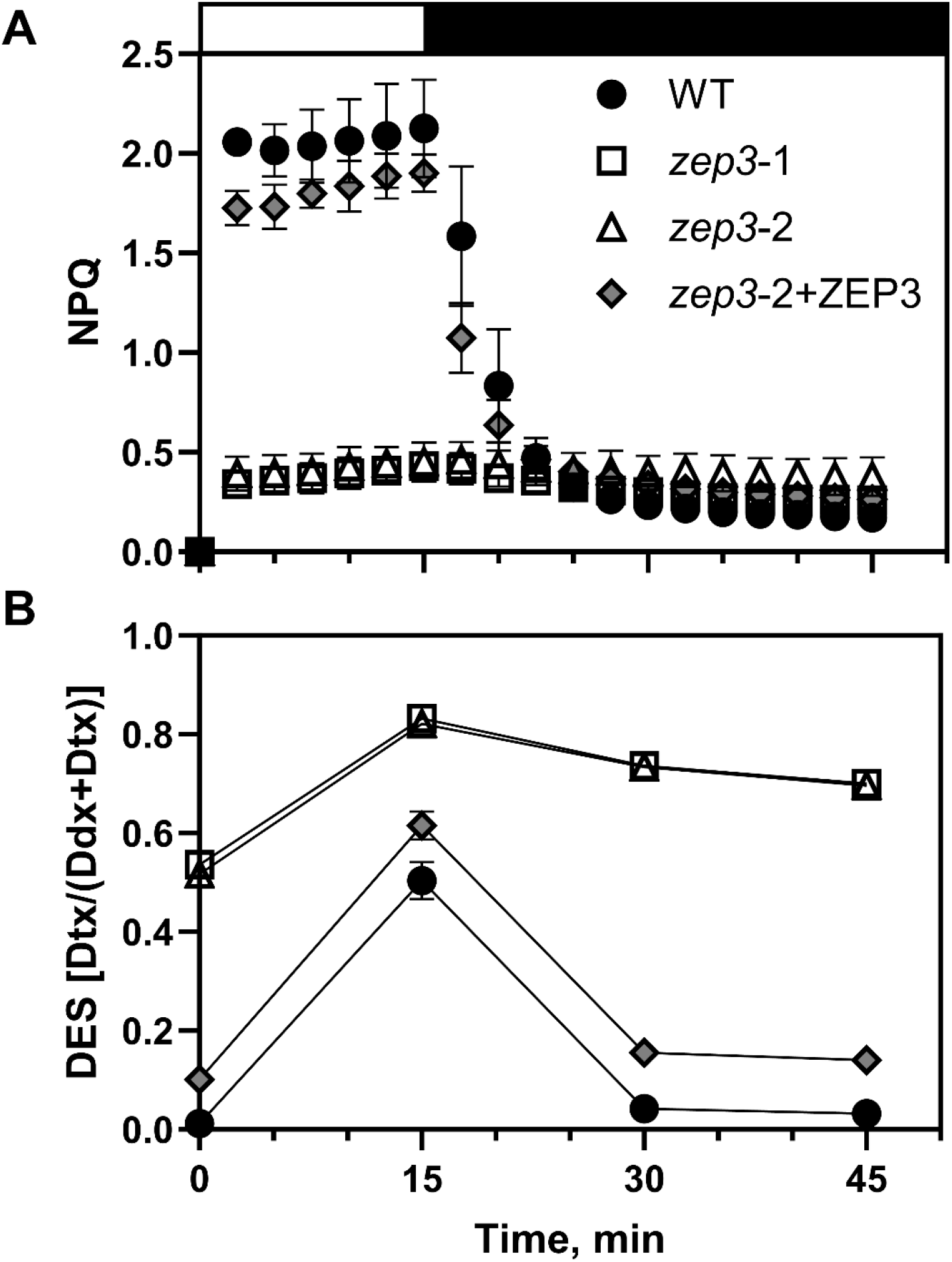
NPQ capacity and de-epoxidation state (DES) of *Phaeodactylum* cultures during an NPQ induction experiment. Cells were exposed to 15 minutes of high light (2000 µmol photons m^−2^ s^−1^, white bar) and 30 minutes of low light (75 µmol photons m^−2^ s^−1^, black bar). NPQ dynamics were measured *ex situ* with a DUAL-PAM fluorometer (A) and DES was estimated via HPLC analyses (B). Dynamics were observed for *Phaeodactylum* WT, two *zep3* mutant strains, and one ZEP3 complemented strain (*zep3*-2 mutant complemented with native ZEP3 gene, *zep3*-2+ZEP3). Cells were grown in sinusoidal light and were collected an hour post-dawn. Points are averages with error bars representing one standard deviation (n=3).

The de-epoxidation state (DES) of a photosynthetic system refers to the proportion of xanthophyll cycle pigments which are de-epoxidated at a given time. In WT diatoms, this is strongly positively correlated with induced NPQ (Lavaud et al., 2002) and is calculated as the amount of diatoxanthin divided by the total amount of diadinoxanthin and diatoxanthin. We therefore also measured the DES of these strains in a parallel NPQ induction experiment. WT started off with a near-zero DES in low light, followed by a large increase within 15 minutes of saturating light (Fig 1B). The DES returned to near-zero between 15 and 30 minutes of low light relaxation (Fig 1B). This suggests a functioning diatoxanthin epoxidase in these conditions. Contrastingly, both mutants began the experiment with a high DES, approximately equal to the peak DES of wild type. The DES signal increased during saturating light and remained high during the low light relaxation (Fig 1B). This suggests a non-functional diatoxanthin epoxidase in the *zep3* mutants. The ZEP3-complemented line exhibited NPQ dynamics comparable to WT and a DES that also was induced and relaxed along with WT, although at a slightly higher baseline than WT (Fig 1). Additional tested ZEP3-complemented lines showed restored inducible NPQ capacity at variable peak levels compared to wild type (Fig S2).

To further inspect the DES dynamics discussed above, we looked at the changing concentrations of diatoxanthin and diadinoxanthin individually. In WT, the epoxidated xanthophyll decreased from an average 383 to 180 mmol diadinoxanthin mol chl *a*^−1^ over the course of the saturating light period, while the de-epoxidated xanthophyll increased from 5 to 183 mmol diatoxanthin mol chl *a*^−1^ in this time frame (Fig 2). This was mirrored by the relative changes at the end of the relaxation period, with a decrease of diatoxanthin concomitant with an increase of diadinoxanthin. In *zep3* lines, we observed a pattern that showed elevated diatoxanthin at the beginning of the experiments, conversion of diadinoxanthin to diatoxanthin, and then little change in epoxidation during the low light phase (Fig 2B). This further supports the role for the ZEP3 gene being required for the de-epoxidation reaction correlating with the reversal of NPQ in low light. Notably, no violaxanthin, antheraxanthin, and zeaxanthin, were observed in our experiments (Fig S3). These are the constituent pigments of the VAZ cycle.

**Fig. 2.**
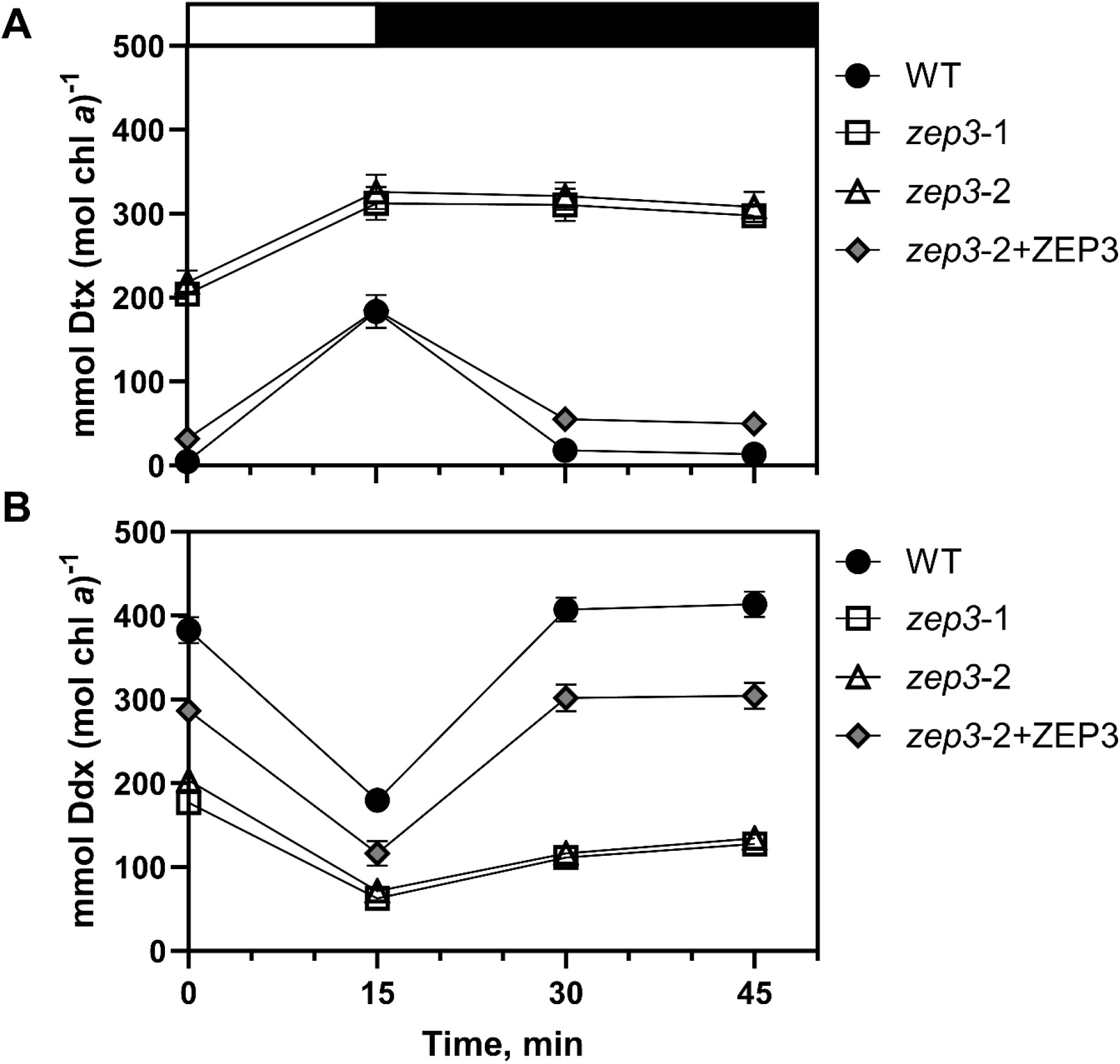
Diadinoxanthin (Ddx) and diatoxanthin (Dtx) dynamics of *Phaeodactylum* cultures during an NPQ induction experiment. Cells were exposed to 15 minutes of high light (2000 µmol photons m^−2^ s^−1^, white bar) and 30 minutes of low light (75µmol photons m^−2^ s^−1^, black bar). Concentrations of Ddx (A) and Dtx (B) were measured via HPLC analyses and normalized to total chlorophyll *a*. Dynamics were observed for *Phaeodactylum* WT, two *zep3* mutant strains, and one ZEP3 complemented strain. Points are averages with error bars representing one standard deviation (n=3).

In a separate series of experiments, we grew the WT and KO strains in constant low light (60 μmol photons m^−2^ s^−1^). We rationalized that the cells would not be exposed to saturating light intensities before measurement (Lavaud et al., 2002) and therefore we could expect the strains to be in a similar photo-physiological state at the start of our assays. This expectation was validated by the comparable values of maximum quantum yield of PSII for WT and the two *zep3 s*trains, which were 0.692, 0.580, and 0.632, respectively (Fig S4D). The WT and two zep3 KO mutants showed rapid induction of NPQ in saturating light, and again little to no reversal of NPQ was observed in the mutant strains compared to the WT (Fig S4A-C).

### Growth of *zep3* is Reduced in Variable Light Regimes

Mutant plants and algae with an inability to regulate NPQ have reduced fitness in excess light and natural light regimes (e.g.(Külheim et al., 2002; Peers et al., 2009). So, we next investigated the role of ZEP3 on fitness of *Phaeodactylum* in several different growth conditions. We used five light regimes, three of which were 12 hr day:12 hr night regimes with variable light during the day. Sinusoidal light (SL) increases from 0 to 2000 μmol photons m^−2^ s^−1^ and then back to darkness in a 12-hour sinusoidal pattern before night (Fig 3A). Intermittent light (IL) comprised repeated intervals of two minutes of darkness and two minutes of 1000 μmol photons m^−2^ s^−1^ light for twelve hours before night (Fig 3A-B). Estuary light (EL) consisted of 8 second spikes from 0 to 2000 μmol photons m^−2^ s^−1^ and back every 30 seconds for twelve hours (Fig 3A-B), modeled from conditions a single cell would experience in a rapidly mixing estuary. In all these light conditions, both *zep3* strains demonstrated lower specific growth rate compared to WT, while the ZEP3-complemented strain recovers growth rates to near wild type-levels (Fig 3C-E). We also deployed two constant 24h light regimes, low (LL) and high (HL) light at 60 and 450 µmol photons m^−2^ s^−1^, respectively (Fig S5A). In these light regimes, *zep3 strains* did not grow slower than wild type (Fig S5B-C). This illustrates the importance of NPQ relaxation for efficient growth in variable light.

**Fig. 3.**
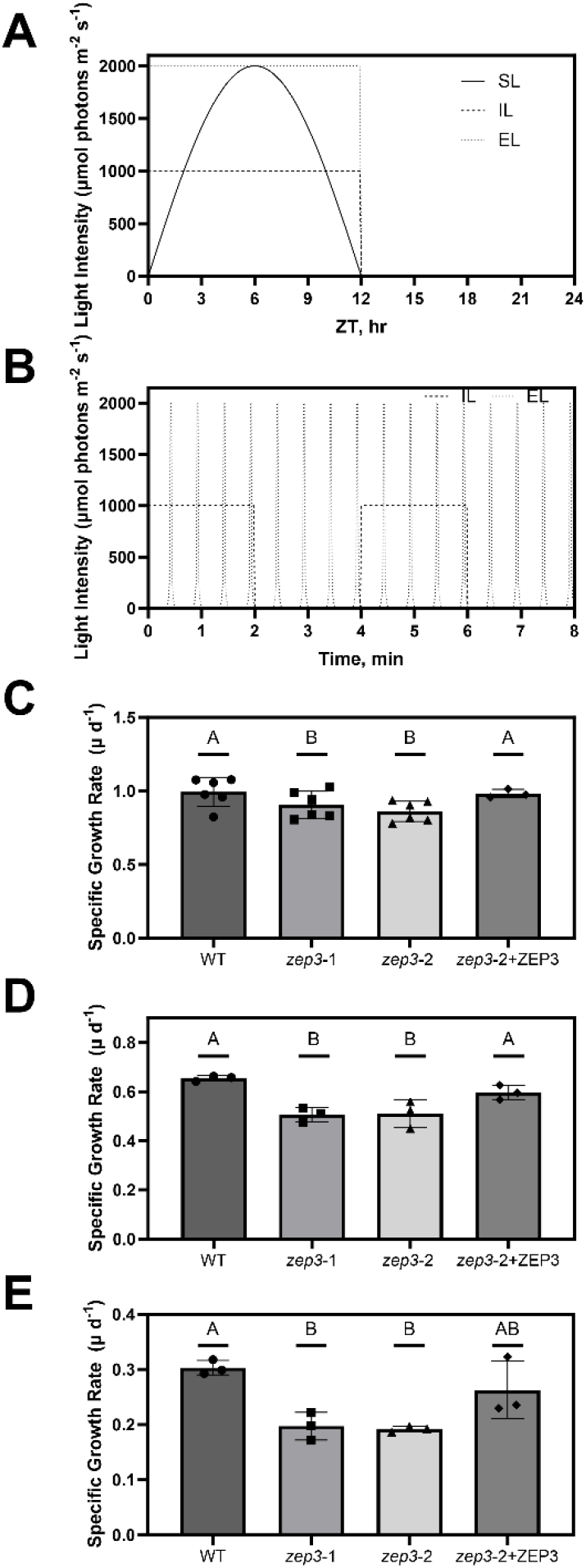
Maximal specific growth rates of *Phaeodactylum* cultures during different 12h D:N variable light regimes. A, Schematic of light regimes utilized in this experiment on a 24 hour time scale, namely sinusoidal light (SL), intermittent light (IL), and estuary light (EL). B, Schematic of the IL and EL regimes on a time scale of several minutes during the day to visualize the finer resolution of light dynamics. C-E, Specific growth rates of cultures grown in SL (C), IL (D), and EL (E) regimes. Growth rates were observed for *Phaeodactylum* WT, two *zep3* mutant strains, and one ZEP3 complemented strain. Bars are averages with points from individual replicate cultures shown and error bars representing one standard deviation (n=6 or 3). Letters represent statistically different groups tested via a repeated measures one-way analysis of variance (RM-ANOVA) with a Tukey’s HSD test.

### Photosynthesis and NPQ of *zep3* are Diminished at Low and Moderate Light Intensities

We sought to understand if reduced growth rates in variable light environments are due to a change in overall photosynthetic activity. Cells were sampled from sinusoidal grown cultures at ±1 hour of solar noon. We performed simultaneous PAM fluorescence and oxygen evolution measurements to quantify changes in photosynthetic performance in a series of increasing light intensities. As expected from prior measurements, the NPQ capacity of *zep3* remains at a low level compared to WT, even at the highest light intensities (Fig 4A). The complemented strain showed an intermediate phenotype. While photosynthetic rates for the WT and complemented strains appeared higher than the KO strain at light intensities above 250 µmol photons m^−2^ s^−1^ (Fig 4B), the calculated maximal rates of photosynthesis (P_max_) were statistically identical between all 3 strains (Table 1).

**Fig. 4.**
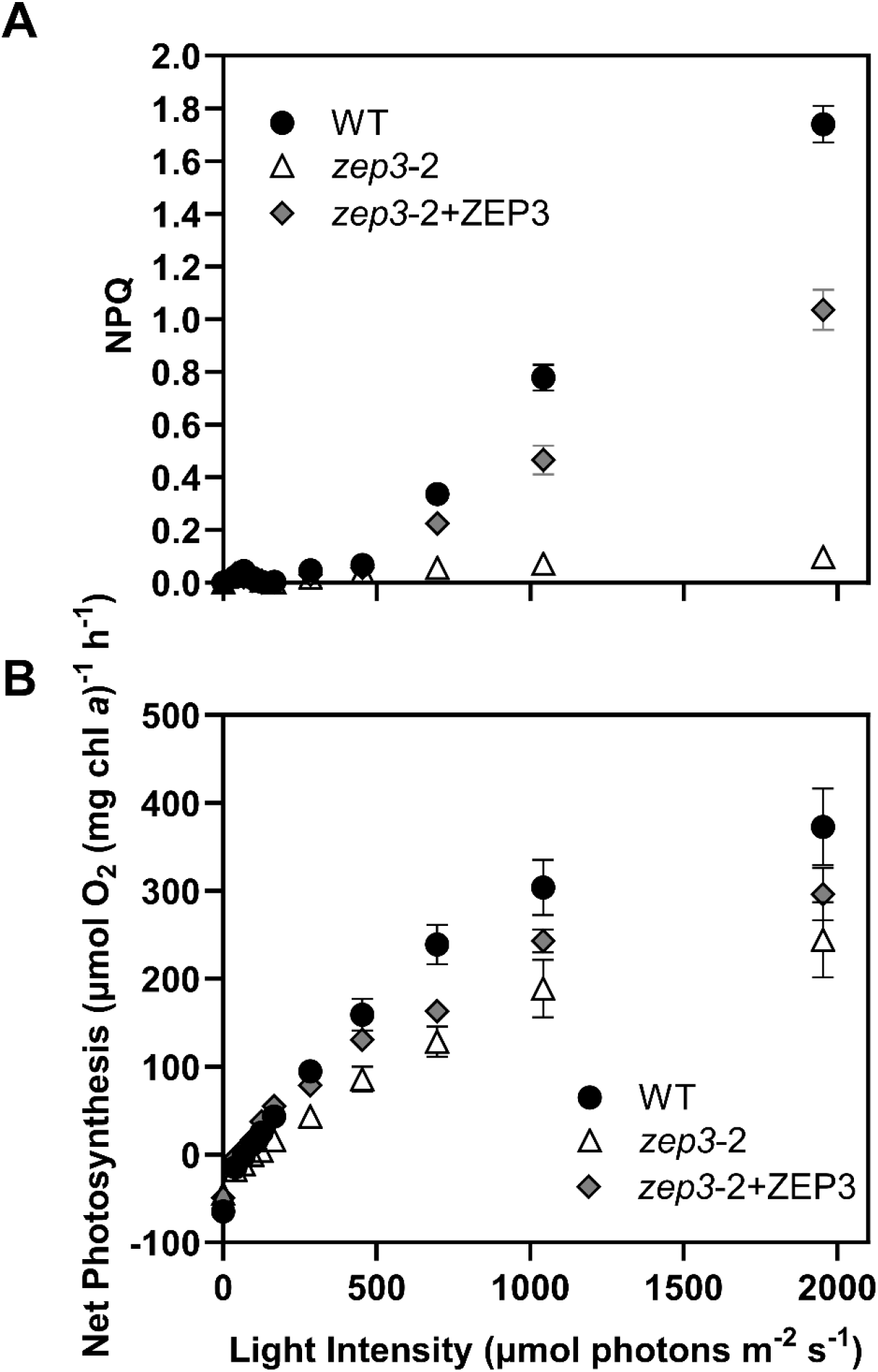
NPQ capacity and net photosynthesis of *Phaeodactylum* cultures during an *ex situ* P-I curve. A, NPQ measured with a DUAL-PAM fluorometer. B, Net photosynthesis derived from gross oxygen evolution rates were measured with a FireSting oxygen probe, corrected for dark respiration rate, and normalized to total chlorophyll *a*. Cells were collected from sinusoidal light grown cultures and samples were taken within 2 hours of solar noon. Dynamics were observed for *Phaeodactylum* WT, one *zep3* mutant strain, and one ZEP3 complemented strain. Points are averages with error bars representing one standard deviation (n=3).

**Table 1.**
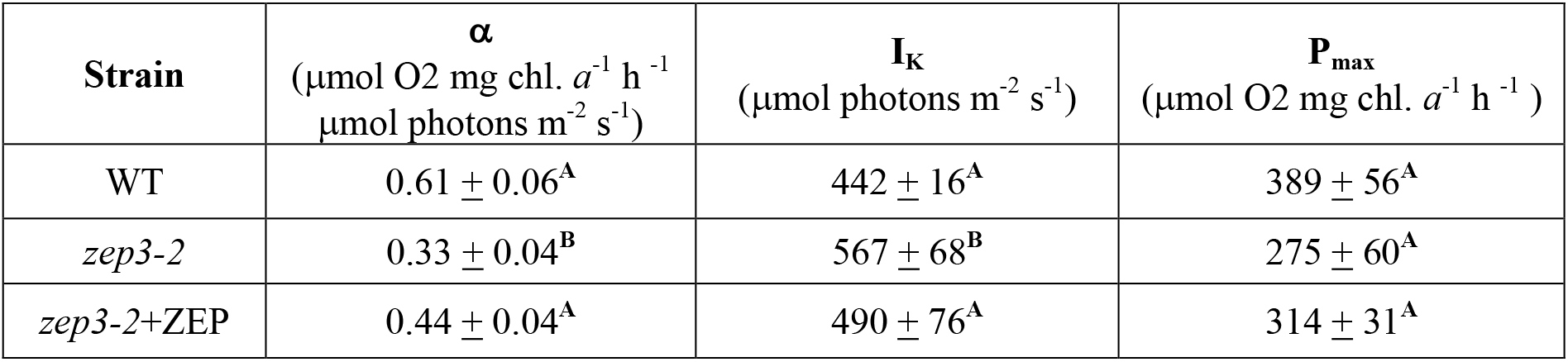
Photosynthetic parameters derived from photosynthesis-irradiance curves. Strains were evaluated for the light limited slope of photosynthesis (α), the irradiance at saturation (I_K_) and maximal rates of photosynthesis (P_max_). Values are averages (n=3) +/− standard deviation. Letters indicate statistical differences between samples (p<0.05). chl = chlorophyll

| Strain | $\alpha$<br>($\mu\text{mol O}_2 \text{ mg chl. } a^{-1} \text{ h}^{-1}$<br>$\mu\text{mol photons m}^{-2} \text{ s}^{-1}$) | $I_K$<br>($\mu\text{mol photons m}^{-2} \text{ s}^{-1}$) | $P_{max}$<br>($\mu\text{mol O}_2 \text{ mg chl. } a^{-1} \text{ h}^{-1}$) |
| --- | --- | --- | --- |
| WT | $0.61 \pm 0.06^A$ | $442 \pm 16^A$ | $389 \pm 56^A$ |
| *zep3-2* | $0.33 \pm 0.04^B$ | $567 \pm 68^B$ | $275 \pm 60^A$ |
| *zep3-2+ZEP* | $0.44 \pm 0.04^A$ | $490 \pm 76^A$ | $314 \pm 31^A$ |

Removal of ZEP3 did affect photosynthetic rates in lower light fluxes. The calculated α parameter represents the efficiency of light harvesting during light-limited photosynthesis. The *zep3-2* mutant had a clearly reduced efficiency compared to the WT and complemented strains (Table 1), suggesting that the “locked” NPQ state reduced overall photosynthetic rate in low light. Correspondingly, *zep3* also had significantly higher irradiance for light saturation (I_K_). These results suggest the lower growth rates observed for the *zep3* mutant in variable light are due to its reduced capacities for light capture in subsaturating light.

### The Capacity to Change Light Harvesting Efficiency in Variable Light is Impaired in *zep3*

It was also of interest to us to investigate how the inability to rapidly regulate NPQ in variable light would affect the light harvesting capacity of *zep3* compared to the WT. We used Fluorescence Induction and Relaxation (FIRe) fluorometry to measure the PSII absorption cross-section of cells sampled from cultures over the course of a sinusoidal day and night. This was done both immediately upon sampling, to approximate the *in situ* absorption cross-section, and following a 20 minute dark relaxation. The absorption cross-section of WT a half hour after dawn was high and had similar values between *in situ* and dark-relaxed states, at 450 and 466 Å^2^ quantum^−1^, respectively (Fig 5). Towards solar noon, both *in situ* and dark-relaxed absorption cross-section values decreased relative to the values following dawn and rose again towards dusk. The *in situ* values changed from 450 Å^2^ quantum^−1^ after dawn to 317 at noon and back to 446 before dusk. Dark relaxation considerably recovered the absorption cross-section value of the WT relative to the values taken immediately after sampling during the day. At noon, the average absorption cross-section value taken upon sampling was 317 Å^2^ quantum^−1^, while the value after dark relaxation was 398. The absorption cross-section decreased somewhat from half an hour before dusk to a half hour before the following dawn, from 446 to 387 Å^2^ quantum^−1^. These trends for WT were mimicked by the *zep3-2*+ZEP3 complemented strain, albeit with slightly lower values across the day.

**Fig. 5.**
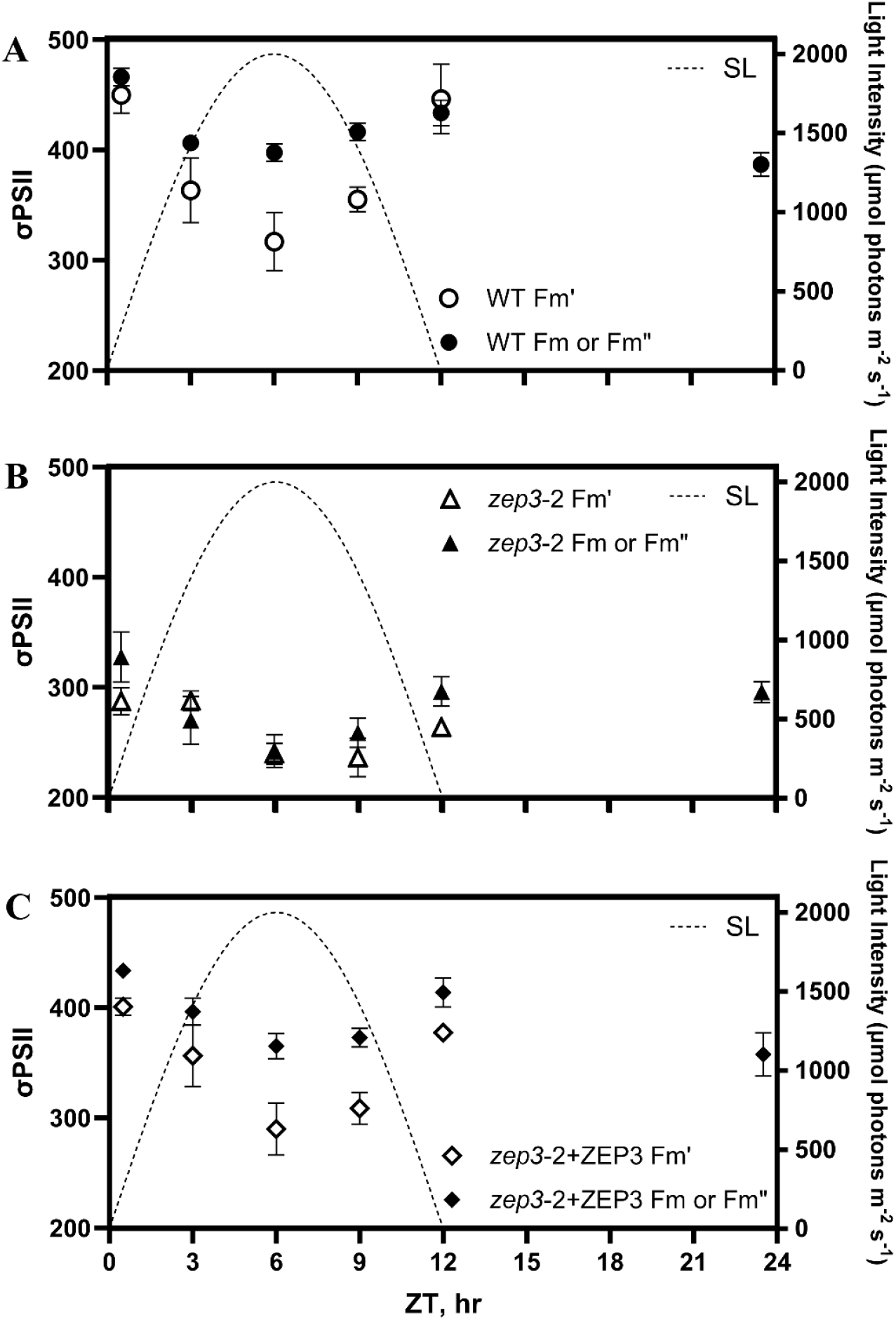
PSII functional absorption cross-sections (σPSII) measured over a sinusoidal day. A-C, Values of σPSII were measured immediately after sampling (white points, Fm′) or following a 20 minute low light relaxation (black points, Fm for predawn sample; Fm′′ for samples collected during the day) with a Satlantic FIRe fluorometer. Measurements were performed for *Phaeodactylum* WT (A), one *zep3* mutant strain (B), and one ZEP3 complemented strain (C). The corresponding sinusoidal light condition is plotted using the right Y-axis. Points are averages with error bars representing one standard deviation (n=3).

In contrast to WT, the absorption cross-section values of *zep3* had less of a dynamic range both over the course of the day and between dark-relaxed and *in situ* states. Values for *zep3* were less than WT across the whole day; for instance, *zep3* showed a dark-relaxed absorption cross-section of 328 Å^2^ quantum^−1^ a half hour after dawn, compared to 466 for WT at that time. The *in situ* value at noon for *zep3* was 242 Å^2^ quantum^−1^, and this compared to 287 after dawn is a much lower change than WT exhibited between these time points, which had a difference of 133 Å^2^ quantum^−1^. The dark-relaxed values from the WT cultures in the middle of the day were much higher than the *in situ* values. This contrasts to the *zep3* cultures where the in situ and dark adapted values were fairly similar. This implies that *zep3* cells are impaired in adjusting their light harvesting capacity in response to natural light variation. This is further supported by the relative lack of change in the absorption cross-section from just before dusk to the end of night, which remained at an average 296 Å^2^ quantum^−1^, in contrast to the larger decrease observed in WT over the night.

### Inducible ZEP3 Expression Causes Dose- and Time-Dependent NPQ Capacity

We then wanted to investigate if we could tune the level of reversible NPQ by adjusting levels of ZEP3. To do this, we modulated the expression of *Pt*ZEP3 using the inducible expression system previously developed in our lab (Kassaw et al., 2022). To that end, we complemented the *zep3-2* mutant by putting the native coding sequence of ZEP3 under the control of an inducible promoter driven by a synthetic β-estradiol-responsive transcription factor (Fig S6A). We obtained several inducible complemented strains confirmed by PCR amplification of the gene sequence as well as the inducible promoter sequence (Fig S6B). The NPQ capacity of these strains was tested with an IMAGING-PAM over 48 hours and over a β-estradiol inducer range of 0-1 μM. An inducible ZEP3 (iZEP3) strain subjected to this inducer range for 24 hours showed a dynamic variation of NPQ capacity of 0.6 to 1.06 from 5 nM to 1 μM β-estradiol (Fig 6A). The lowest inducer concentrations of 0.1 to 1 nM did not result in a detectable change in NPQ relative to the vehicle control of ethanol (NPQ ∼0.38). In comparison, wild type displayed relatively consistent NPQ capacity in the tested range of β-estradiol (range = 0.99 to 1.14). This is comparable to that of the iZEP3 strain at the highest used β-estradiol concentration (Fig 6A). Induction of NPQ in the iZEP3 strain was also time-dependent as high NPQ capacity at the highest β-estradiol concentration was observed 12 hours after induction and remained high through 48 hours (Fig 6B). Notably, some increase in NPQ capacity was already observed after 4 hours of induction (Fig 6B). The broader dynamic range of NPQ capacity in the iZEP3 strain can be seen across both time and dosage, reiterating that high NPQ is reached at 12 hours and dynamic range is observed from 5 nM to 1 μM β-estradiol (Fig S7).

**Fig. 6.**
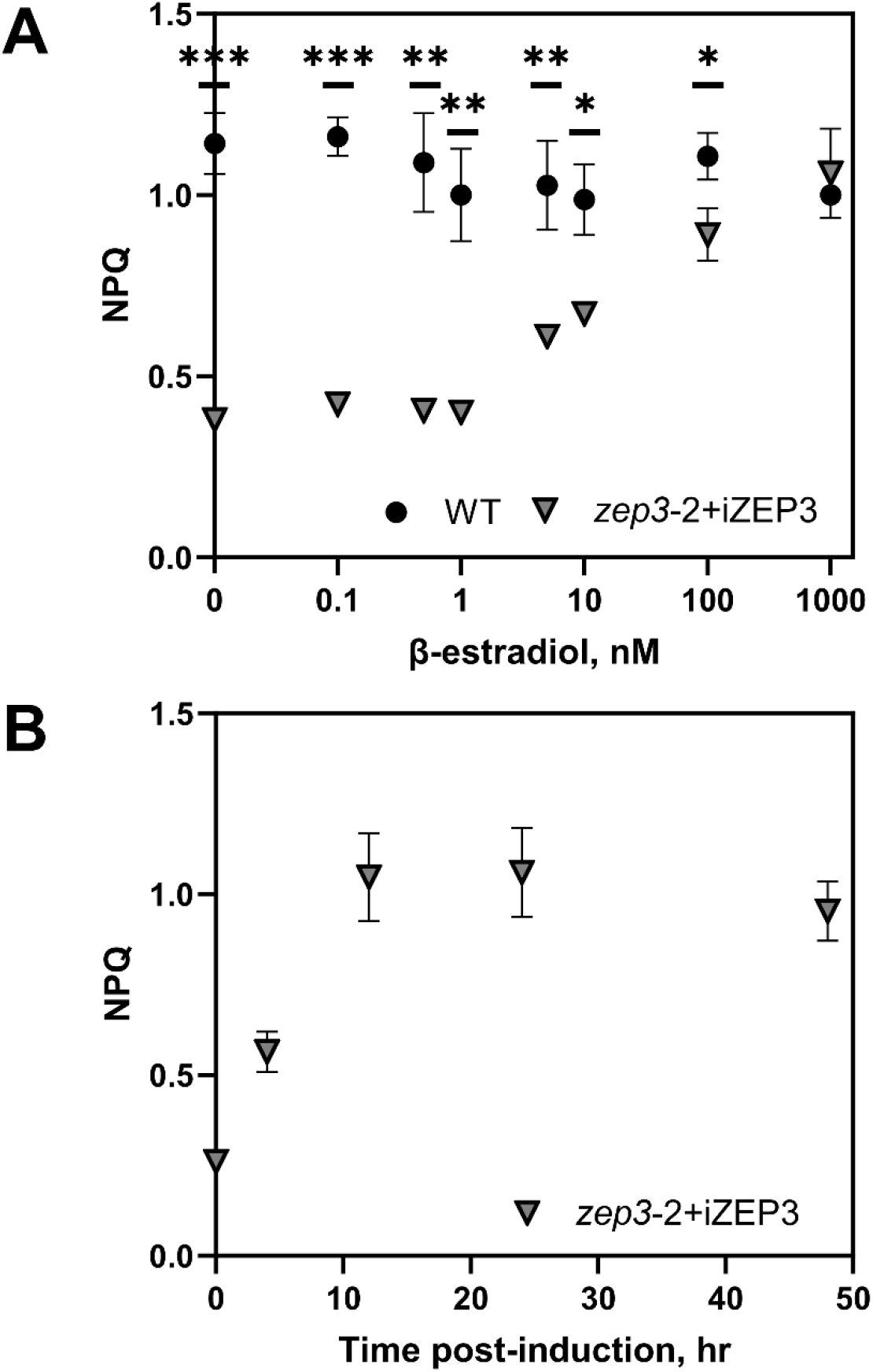
Dose- and time-dependent tuning of NPQ capacity via chemical induction of ZEP3 expression. An inducible ZEP3 *Phaeodactylum* strain (*zep3*-2+iZEP3) was created via complementation of the *zep3*-2 mutant with a construct containing the native ZEP3 gene under the control of a β-estradiol-inducible synthetic promoter. A, The NPQ of the iZEP3 strain and WT was measured with an IMAGING-PAM at different β-estradiol concentrations after 24 hours. B, The NPQ of the iZEP3 strain was measured over 48 hours at 1000 nM β-estradiol. Cells were incubated and assayed in 24 well plates with 2 mL of f/2 medium to which was added 2 µL of β-estradiol at the needed concentration or 2 µL of ethanol as a vehicle control. Points are averages with error bars representing one standard deviation (n=3). Asterisks represent statistically significant differences between WT and the iZEP3 strain, tested via unpaired t-tests at each timepoint with Holm-Šídák’s multiple comparison method (*: p<0.05, **: p<0.005, ***: p<0.0005).

## Discussion

NPQ is a collective term for the quenching of chlorophyll fluorescence. This is achieved via an array of molecular mechanisms to dissipate absorbed photons upstream of the photosystem reaction centers. These mechanisms have been termed: energy-dependent (qE, Wraight and Crofts, 1970), LCNP-SOQ-ROQH1-dependent (qH, Malnoë et al., 2018), photoinhibitory (qI, Krause, 1988), chloroplast avoidance movement (qM, Dall’Osto et al., 2014), state transition (qT, Bonaventura and Myers, 1969), and zeaxanthin-dependent quenching (qZ, Nilkens et al., 2010). Other factors, such as electrochemical gradients (Cardol et al., 2010), ion homeostasis (Kunz et al., 2014), and UV (Allorent et al., 2016) contribute to NPQ capacities and kinetics. Here we will focus on the two major mechanisms, qE and qZ, as these have also been identified to play significant photoprotective roles in algae and particularly in diatoms.

The well characterized qE response in plants is driven by an accumulation of protons in the thylakoid lumen. The acidification results in changes in the supercomplex structure of PSII, which favours energy transfer to lutein in LHCII and the dissipation of energy as heat (Ruban et al., 2007). Luminal acidification causes the LHC family protein PsbS to bind to major and minor antennae complexes to accelerate this process (Sacharz et al., 2017). Green algae have been shown to possess a similar cascade of events. Green algae (Peers et al., 2009), mosses (Alboresi et al., 2010), and some streptophytes (Gerotto and Morosinotto, 2013) possess LHCSR as well as PsbS. The more closely related the green alga is to a streptophyte, the more significant a role PsbS plays in NPQ, and conversely LHCSR plays a reduced role.

qZ is an NPQ component directly proportional to the “light-dissipative” carotenoid present in the light harvesting antennae. The induction of qZ is significantly affected by qE, but its reversal is ΔpH independent. In higher plants and green algae, this carotenoid is zeaxanthin. The VAZ cycle is described in the introduction of this manuscript.

Diatoms, hacrobians, and alveolate algae possess LHCX and LHCX-like proteins which are analogous to LHCSR. LHCX/LHCX-like proteins accumulate in stress conditions like LHCSR and PsbS (Bailleul et al., 2010), but the mode of action which increases thermal energy dissipation is not yet clear (Giovagnetti and Ruban, 2017). The VAZ cycle can also be observed in diatoms (Lohr and Wilhelm, 1999), but diatoms, haptophytes, and dinophytes also utilize an independent one-step xanthophyll cycle. The Ddx:Dtx cycle has also been suggested as solely responsible for NPQ in diatoms (Blommaert et al., 2021). A unique feature of this cycle, presumably due to chlororespiration maintaining low pH, is a residual pool of diatoxanthin that exists in darkness. Here we aimed to identify a gene responsible for the Ddx/Dtx cycling in diatoms and investigate its importance in NPQ dynamics.

### ZEP3 is required for the epoxidation of diatoxanthin and NPQ relaxation

We focused our physiological characterization on two *zep3* knockout (KO) strains along with a single strain that had been complemented with the native ZEP3 gene. Both the WT and ZEP3 complemented strains showed rapidly inducible and reversible NPQ when challenged with excess light (Fig. 1). The mutant strains had considerably lower inducible NPQ that did not revert following a shift to low light. Correspondingly, the WT and the complemented strain showed the expected change of the xanthophyll cycle pigments – an increase in Dtx content in excess light and its reversal to Ddx in low light. The *zep3* mutants started the excess light treatment with elevated Dtx, followed by additional conversion of Ddx to Dtx during excess light and very little re-epoxidation during low light recovery (Fig. 1, Fig 2, Fig. S3). This result led us to assume that ZEP3 is likely the Dtx epoxidase enzyme responsible for reversing NPQ back to the light harvesting state. This illuminates the diversification of function for the ZEP family in Ochrophytes as ZEP1 is known as an integral part of the fucoxanthin biosynthesis pathway (Bai et al., 2022). We highlight that we did not measure the catalytic activity of the ZEP3 gene product which will be required to irrevocably assign this protein as the Dtx epoxidase. The role of ZEP2 was not investigated in our work.

### ZEP3 is required for maximum growth rates in variable light regimes

Traditionally, static light regimes with 24h day or square wave 12h day:night cycles, have been used in laboratories to study photosynthesis. Increasingly, intermittent, variable or more natural light regimes are being employed to study the contribution of individual proteins and pathways involved in excess energy dissipation pathways such as alternative electron transport (Allahverdiyeva et al., 2013; Andersson et al., 2019; Nawrocki et al., 2019). We initially characterized the growth of the *zep3*, complemented and WT strains in constant light. Low light (LL) and high light (HL) culturing conditions revealed no changes in relative growth rates between the three groups (Fig S5). We hypothesized that very little NPQ is induced in these conditions due to physiological acclimation to the uniform light flux (Jallet et al., 2016).

However, NPQ is expected to play an important role in conditions where light fluxes exceed the ability for cells to utilize absorbed light energy for photosynthesis (Murchie and Ruban, 2020). We designed 3 different conditions to assay the effect of variable light on fitness (growth rate). The first condition was a sinusoidal light (SL) condition, mimicking conditions in the upper water column on a sunny day with no mixing ((Ware et al., 2020); see Fig. 3 for graphical comparison of light conditions). The second condition used an intermittent light series (IL) where cells move in and out of saturating light in a rapid fluctuation. The IL light series has been shown to induce massive NPQ capacity in diatoms (Ruban et al., 2004). The third condition is an estimate of mixing in an estuarine environment (EL) where cells quickly mix between the surface and depth. We used light attenuation and water column mixing rate data to recreate a light environment that could occur in an estuary. We found that each of these conditions resulted in a lower growth rate for the *zep3* mutants compared to the WT (Fig. 3). The complemented strain recovered to WT growth rates in SL and IL light conditions. But we note the complemented strain was not statistically different from either WT or *zep3* mutants in the EL condition, though a similar trend to the other conditions was apparent. These results offer two major takeaways. Firstly, ZEP3 and the conversion of Dtx back to Ddx is critical for maximum growth rates in dynamically changing light environments, such as those found in natural environments. Secondly, simulating natural light environments can elucidate key processes that are required for maximal growth rates of algae. While there are clear demonstrations that algae with the inability to induce NPQ have reduced fitness in excess light or rapidly changing light due to increased oxidative damage (Peers et al., 2009; Bailleul et al., 2010), our results demonstrate the capacity to reverse NPQ is also required for maximal growth in variable light.

### ZEP3 is essential for high rates of photosynthesis

To test whether the decreases observed in cellular fitness in variable light were the result of impaired photosynthetic performance, we assayed photosynthesis via simultaneous PAM fluorescence and oxygen evolution measurements. Photosynthesis-irradiance curves have long been employed to estimate the efficiency of light harvesting (MacIntyre et al., 2002). We found WT NPQ was induced at light fluxes above 500 μmol photons m^−2^ s^−1^ (Fig. 4). As expected, the *zep3-2* mutant showed little change in NPQ capacity throughout the experiment while the complemented strain showed an intermediate phenotype. The *zep3-2* mutant showed reduced levels of oxygen evolution compared to the WT as indicated in Table 1. Therefore, ZEP3 is required to maximise the efficiency of light harvesting and being locked in an NPQ state (high Dtx) results in lower rates of photosynthesis. In higher plants, the *npq2* mutant, devoid of the zeaxanthin epoxidase, has lower quantum yields of PSII (ΦPSII) at low light intensities but not at high light, compared to WT (Ware et al., 2016). Our work demonstrates that *zep3* mutants are impaired in recovery of ΦPSII following a high light challenge (Fig. S4D), which provides further justification for the lower cellular fitness (growth rate) we observed in variable light.

### Natural light regimes require dynamic changes in the cross section of PSII

It is assumed that NPQ processes exert a strong influence on the variable fluorescence observed in natural phytoplankton communities, with clear reductions of fluorescence yields associated with excess light (Gorbunov and Falkowski, 2022). Our *ex situ* estimations of photosynthetic performance for cells grown in sinusoidal light suggested that *zep3* had reduced light harvesting efficiencies (Fig. 4), which was likely influenced by cells being locked in an induced NPQ state. NPQ in *Phaeodactylum* has been correlated with a decrease in the functional absorption cross section of PSII (σPSII, (Giovagnetti and Ruban, 2017; Buck et al., 2019). The σPSII has also been demonstrated to vary significantly during the course of sinusoidal light cultivation, presumably due to the induction/reversal of NPQ (Jallet et al., 2016). We probed whether the variations in the σPSII were influenced by ZEP3. Cultures were assayed within 30s of removal from *in situ* SL conditions to estimate the σPSII *in vivo*. Estimations of the σPSII suggested that ZEP3 is required *in situ* to alter the σPSII (Fig. 4). WT and ZEP3-Comp cells had a σPSII almost twice as large as ZEP3 cells at dawn and dusk. At solar noon, in conditions of 2000 μmol photons m^−2^ s^−1^ there was only a small difference in the σPSII of ZEP3 to the WT *in situ*. In lower light environments after solar noon, ZEP3 cells have fewer antenna proteins delivering excitation energy to PSII, which is supported by the reduced light harvesting efficiency inferred from oxygen evolution measurements (Fig 4). Reduced rates of oxygen evolution are therefore likely a contributing factor to the reduced fitness of ZEP3 KOs. One of the reasons for reduced photosynthetic performance is the irreversible accumulation of diatoxanthin (Fig. 2A), which reduces the light harvesting capacity of *zep3* by irreversibly truncating σPSII. Overall, this illustrates the importance of reversing from NPQ to a light-harvesting state in a natural environment.

### Synthetic circuits can be utilised to control ZEP3 expression in a time and dose-dependent manner

NPQ is a target for engineering photosynthetic efficiency (Murchie and Ruban, 2020). The concept associated with this engineering target is that dense algal cultures or high-density canopies lead to highly variable light environments that change faster than native NPQ mechanisms can turn on and off. So, fast transitions from excess light to low light may result in lower light harvesting capacity due to the persistence of NPQ. Indeed, work in cyanobacteria and plants has shown that the deletion of NPQ capacity (Peers 2015) or tuning of NPQ induction and relaxation can lead to increased biomass accumulation in agricultural situations or dense algal cultures (Kromdijk et al., 2016; De Souza et al., 2022; Perin et al., 2023). We investigated if the level of NPQ could be tuned in *Phaeodactylum* using a synthetic promoter system. We modified our inducible expression system (Kassaw et al., 2022) to incorporate the native ZEP3 coding sequence under the control of a β-estradiol-dependent promoter into *zep3* cells (*zep3*-iZEP3; Fig S6). Within 4 h of induction, the phenotype of reversible NPQ could be observed in the 100-1,000 nM β-estradiol treatments. Additionally, β-estradiol concentrations as low as 5 nM partially restored NPQ reversal within 12 h. This shows NPQ can be controlled like a rheostat in a time and dose dependent manner. Employing synthetic circuits to control the expression of photosynthetic and photoprotective enzymes could allow algal cultivators to adjust the photosynthetic performance of algae in near real-time depending on the growth phase of cultivated algae. Future studies could investigate if biomass or bioproduct formation could be enhanced in mass cultures of algae using synthetic control of NPQ capacity.

## Conclusion

Diatoms contend with mixing in the water column and resulting fluctuations in light levels, which provides environmental selection for efficient NPQ. In several ways, diatoms have less complicated mechanisms of NPQ than land plant and green algal counterparts. Diatoms do not exhibit state transitions (qT) and therefore rely heavily on xanthophyll cycling (qE) for NPQ on shorter time scales. While in plants, xanthophyll cycling involves the generation of two de-epoxidated xanthophylls, antheraxanthin and zeaxanthin, diatoms only remove one epoxide ring from Ddx to form Dtx for photoprotection. Our work demonstrates that a member of the *Phaeodactylum* ZEP family, ZEP3, likely functions as the diatoxanthin epoxidase involved in NPQ. This enzyme is needed for rapid reversal of NPQ, cellular fitness in variable light environments, and normal photosynthetic rates and photo-physiological parameters under high light. We also demonstrate that inducible expression of ZEP3 leads to variable levels of NPQ. This has potential ramifications for engineering photosynthetic efficiency in industrial culture of diatoms in different light regimes of interest.

## Materials and methods

### Growth conditions

*Phaeodactylum tricornutum* CCAP 1055/1 was grown axenically in Instant Ocean artificial seawater medium (35.95‰ salinity), supplemented with 0.03% Proline A and Proline B micronutrient stocks. Cultures were maintained in exponential growth phase (approx. 5×10^4^ – 2×10^6^ cells ml^−1^) throughout this study. Cultures were grown under constant shaking (VWR shaker model 3500, shaker setting 4), 18^°^C, and ambient air. Light intensities were measured using the Walz ULM-500 universal light meter. Five different light regimes were used to assay fitness in this study. Two are 24h constant light regimes at high (HL; 450 μmol photons m^−2^ s^−1^) or low light (LL; 60 μmol photons m^−2^ s^−1^). The remaining three are 12h D:N cycles with variable light intensity. The sinusoidal light (SL) regime follows this formula:

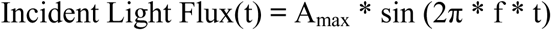

Where t = h after dawn, A_max_ = 2000 μmol photons m^−2^ s^−1^, f = 1/(DayLength[h])=1/24. The light changes in intervals of 5 minutes. The intermittent light (IL) regime followed a square wave pattern (day: 120s 1000 μmol photons m^−2^ s^−1^, 120s darkness, etc.). The estuary light (EL) regime was based around environmental conditions measured in two different estuaries. Water turbidity measurements in the Chesapeake Bay suggested less than 5% of incident light is available below 1 meter depth (Carter and Rybicki, 1990). Measurements of water movement in the Columbia River estuary suggested an average mixing depth of 7m and an average mixing efficiency of 0.36. Combining these two studies, we then simplified the complexity of mixing and assumed that cells would spend 1/7^th^ of their time in the light, mixing up into full sunlight and then down again into darkness for the remainder of the 6/7^th^s of the time. The EL was programmed as a series of short spikes (day: 1s intervals of 37, 99, 269, 2s of 2000 μmol photons m^−2^ s^−1^, followed by a mirrored descending pattern and then 22s of darkness etc.).

### ZEP3 identification

Published nucleotide sequences of *Phaeodactylum tricornutum* from the Ensembl protists genome browser was used for identifying target genes. The *Arabidopsis thaliana* Zeaxanthin epoxidase (ZEP) gene (TAIR locus AT5G67030) was used as the query. The ZEP3 gene is annotated as Phatr3_J10970 in the *Phaeodactylum tricornutum* published genome (ASM15095v2).

### Generating HDR mutants of ZEP3

CRISPR-Cas9 mediated homology directed repair (HDR) was used to generate successful knock-outs of the ZEP3 gene in *Phaeodactylum tricornutum* (Fig S1). Two independent guide RNA sequences (Supplemental File 1 for all sequences and primer details) were designed utilising the motif [5’-N20-NGG-3’] against the *P. tricornutum* genome. Oligo design, annealing and synthesis were performed as previously described (Bai et al., 2022) using BsaI-HF®v2 (NEB) digested vector pKS diaCas9_sgRNA (Addgene #74923). The homology donor plasmid was constructed via Gibson assembly (ThermoFisher GeneArt; Gibson Assembly HiFi master Mix #A46628) by fusing 1 kb 5’ homologous arm, ble cassette and 1 kb 3’ homologous arm onto the BamHI (NEB) digested backbone of pUC57. Homology arm sequences and oligos sequences are detailed in Table S1. Linearization of the homology donor and pKS diaCas9_sgRNA plasmids were performed by KpnI-HF® and NdeI (NEB) restriction enzymes respectively. For each electroporation, 100 ml of mid exponential growth phase cells was harvested at 4^°^C, 2500 ×*g*, 10 min. The supernatant was carefully discarded prior to four washes of the pellet in ice cold filtered 0.375 M sorbitol. Cells were resuspended in a final 100 µL of ice-cold 0.375 M sorbitol. 40 µg of heat denatured salmon sperm (1 min 100^°^C) and 2 µg of linearised homology donor and pKS diaCas9_sgRNA plasmids were added to cells and incubated on ice for 10 min. This mixture was transferred to a 2 mm ice cold electroporation cuvette (Bio-Rad). Transformation of WT *P. tricornutum* was performed via electroporation (Bio-Rad GenePulser Xcell, exponential decay, 0.5 kV, 25 μF, 400 Ω). Cells were immediately mixed with 10 mL of 18^°^C f/2 medium in a 15 mL conical tube. Cells were placed under 30 μmol photons m^−2^ s^−1^ for 24 h for transformation recovery. Cells were then spread onto selection plates (f/2 + 100 μg/mL zeocin 1% agar) in 24 h 80 μmol photons m^−2^ s^−1^ light conditions.

### Complementation of knockout mutants with native ZEP3 gene

For complementation of knockouts, Phusion^®^ HF DNA polymerase (NEB, USA) mediated PCR amplification of the ZEP3 gene included all exon, intron, promotor and terminator regions of the WT native ZEP3 (primers 4/21 SI Table 1). PCR amplification of a Blasticidin S deaminase (BSD) cassette was performed as previously described (Bai et al., 2022). 2 μg of the BSD and ZEP3 amplicon sequences were used in electroporation, as described above. Selection was performed by plating transformants on f/2 + 10 μg/mL blasticidin S HCl (ThermoFisher Scientific) 1% agar plates under 24 h 80 μmol photons m^−2^ s^−1^ light conditions.

### β-estradiol inducible ZEP3 expression

First, overlapping PCR was employed to generate a ZEP3-NosT fragment. ZEP3 and NosT sequences were PCR amplified separately using primer pairs 9/10 and 11/12 respectively. Overlapping PCR products were generated using primers 9 and 12 flanked by AvrII and SbfI restriction sites (SI Table 1). Phusion^®^ HF DNA polymerase (NEB, USA) was used for all amplification steps. The ZEP3-NosT PCR product was sub-cloned into the pJET vector using the Blunt-End Cloning Protocol (ThermoFisher K1232) and the PCR product of primers 13 and 14 (SI Table 1) was sequence verified. The ZEP3-NosT fragment was then cloned into the TKK031 vector (Kassaw et al., 2022) by digesting the plasmid with AvrII, SbfI and BamHI. The pJET_ZEP3-NosT plasmid was digested with AvrII and SbfI restriction enzymes (NEB). These were then ligated with T4 DNA ligase (NEB) according to manufacturer instructions. For double selection in our knockout strain background, we needed to change the antibiotic resistance gene. For this, we transferred the inducible ZEP3 expression cassettes from TKK031 to a PtPUC3 background containing a blasticidin resistance cassette (Kassaw et al. 2022). BstxI and SbfI unique restriction sites were used for this purpose. Transformation and screening of strains were performed as described above.

### Genotyping

Cell biomass (∼10^6^ cells) was heated at 85^°^C in 50 μL TE buffer (50 mM Tris HCl, 10 mM EDTA, pH 8.0) for 10 min. Samples were cooled on ice, before adding 50 μL of TE buffer and vortexed. Samples were centrifuged at 10,000 ×*g* at 18^°^C for 10 min. The supernatant was carefully removed and frozen at −20^°^C until required for PCR analysis. 1 µL of extract was used as a DNA template for 1× Gotaq^®^ Green Master Mix (Promega) 25 μL volume PCRs. Full primer sequences listed in SI Table 1 were used for sequence verification. All gel electrophoresis experiments were performed in 1% agarose, 0.01% gel red, TAE buffer (40 mM Tris, 20 mM acetic acid, 1 mM EDTA).

### Cell density and growth measurements

A BD Accuri C6 flow cytometer gated for chlorophyll autofluorescence (excitation 488 nm, emission 670 nm) was used to measure culture cell densities. 1 mL of cells was passed through a 30 μm filter prior to sampling. Flow cytometric cell counting was performed at a flow rate of 36 μL min^−1^ for 40 secs, with a minimum number of 1,000 cells measured for each sampling timepoint. Cultures grown under a 12 h D:N photoperiod were measured within +/− 1 h of solar noon each day.

### Chlorophyll extraction and quantification

Cells were collected and TWEEN 20 (0.01% final concentration, Sigma-Aldrich, p7949) was added. Samples were harvested after centrifugated at 10,000 ×*g* for 10 min at 18^°^C and discarding the supernatant. Chlorophylls were extracted in 100% HPLC grade methanol for 10 min in the dark. Cell debris was removed via centrifugation at 20,000 ×*g* for 10 min at 18°C (Porra, 2002). Concentrations of chlorophyll *a* and *c* were calculated using established absorbance and extinction coefficients (Ritchie, 2008).

### HPLC

Cultures were grown under the SL regime (see section “Growth Conditions”). At 1 h post dawn, cultures were collected and transferred to low light (75 μmol photons m^−2^ s^−1^) for 60 min to relax NPQ and ensure maximal diatoxanthin epoxidation. Exponentially growing cultures (average 1.04×10^6^ cells/ml) were sampled at a volume corresponding to a total 7.5 μg chlorophyll *a*. Cells were then exposed to 2,000 μmol photons m^−2^ s^−1^ for 15 min before harvesting the same volume. After placement at 70 μmol photons m^−2^ s^−1^ for the 15 minute and the 30 minute timepoints, cells were harvested again. Cells were collected via gentle vacuum onto a GF/6 filter. Cells were harvested in the current light condition before being placed in a cryovial and frozen in liquid nitrogen within 15 s of sampling and then lyophilized. Pigments were extracted from the filters and analysed by HPLC using HPLC system II as described in Bai et al. (2022).

### PAM NPQ state estimations corresponding to HPLC analysis

Cultures were grown under the SL regime. At 1 h post dawn, cultures were collected and transferred to low light (75 μmol photons m^−2^ s^−1^) for 60 min to relax NPQ and ensure maximal diatoxanthin epoxidation. Exponentially growing cultures were sampled at a volume corresponding to a total 2.5 μg chlorophyll *a* and collected on artificial leaf disks (glass fibre prefilters) pre-soaked in 18^°^C f/2 media. F_v_/F_m_ was measured (saturating pulse (SP) of 10,000 μmol photons m^−2^ s^−1^, duration 0.6 s). Cells were illuminated with 1994 μmol photons m^−2^ s^−1^ for 15 min (SP applied every 150s). The actinic light level was reduced to 72 μmol photons m^−2^ s^−1^ for 30 min with an SP applied every 150 s.

### PAM NPQ state estimations for complemented strain screening

Cultures were grown under the HL regime. Cells were sampled at a volume corresponding to a total 2.5 μg chlorophyll *a* and then placed in 30 μmol photons m^−2^ s^−1^ white light conditions for 20 min prior to screening. They were then collected on artificial leaf disks and subjected to the same light regime as described above (see “PAM NPQ state estimations corresponding to HPLC analysis” section).

### PAM NPQ state estimations for low light measurements

Cultures were also grown under the LL regime. Cells were sampled at a volume corresponding to a total 7.5 μg chlorophyll *a* and then collected on artificial leaf disks pre-soaked in 18^°^C f/2 media. F_v_/F_m_ was measured (SP of 10,000 μmol photons m^−2^ s^−1^, duration 0.6 s). Cells were illuminated with 1996 μmol photons m−2 s −1 for 10 min (SP applied every 60s). The actinic light level was reduced to 79 μmol photons m^−2^ s^−1^ for 10 min with an SP applied every 120 s.

### Simultaneous PAM fluorometry and oxygen evolution measurements

The equipment set-up for simultaneous measurements was the same as previously described (Jallet et al., 2016). All sampling and measurements were performed at 18^°^C. Samples collected ± 2 h of solar noon. Chlorophyll quantification was performed as described above. Samples were collected via centrifugation (18°C, 2500 ×*g*, 10 min, with a final concentration of 0.01% Tween 20). The supernatant was discarded and cells were resuspended in pre-cooled air saturated f/2 media. 2 mL of cells at a concentration of 4 μg chl *a* mL^−1^ were used for measurements. The stopper was lowered to purge air from the sample, leaving a 1.4 mL volume for measurements. Cells were exposed to darkness for 5 min to measure dark respiration rates. 15 minutes of blue actinic light (79 μmol photons m^−2^ s^−1^, BL int 5) was used to relax NPQ. Immediately following the relaxation, the actinic light was turned off and F_v_/F_m_ was measured (SP of 10,000 μmol photons m^−2^ s^−1^, duration 0.6s). Cells were then exposed to 2 minute steps of increasing actinic light intensities of 40, 67, 95, 125, 166, 284, 453, 697, 1042, and 1953 μmol photons m^−2^ s^−1^, with an SP applied at the end of each step. Saturating pulses were delivered to derive F_o_, F, and F_m_, with these parameters used to calculate NPQ and ΦPSII according to published equations (Maxwell and Johnson, 2000). Gross oxygen evolution capacities were normalised to chl *a* concentrations after offsetting the dark respiration rates. The maximal rate of photosynthesis (P_max_), minimum light at saturation (I_K_) and the quantum efficiency of photosynthesis (α) were calculated as described in Ware et al. 2020.

### *In situ* estimations of functional absorption cross sections of PSII

Exponentially growing cultures were re-established at 50,000 cells mL^−1^ at solar noon the day preceding measurements. At the specified timepoints, 1 mL of culture was transferred to a 1 cm quartz cuvette. The functional absorption cross section of PSII (σ_PSII_; Å^2^ quantum^−1^) was measured within 5 s of sampling using a Satlantic FIRe Fluorometer (excitation at 450 nm, emission at 678 nm). A 1.1 mL sample was collected from the same culture and placed under 30 μmol photons m^−2^ s^−1^ white light for 20 min. σ_PSII_ from a 1 mL subsample of this was measured after the low light relaxation to estimate the Fm” state. Fireworx software (https://sourceforge.net/projects/fireworx/) was employed to calculate σ_PSII_ from the raw fluorescence data. Sampling timepoints correspond to 0.5 h, 3 h, 6 h, 9 h, 11.5 h and 23.5 h after dawn during a 12 h day:night sinusoidal light regime (2,000 μmol photons m^−2^ s^−1^ maximum).

### Inducible ZEP3 expression NPQ relaxation dynamics

Cultures were maintained under 450 μmol photons m^−2^ s^−1^, 20^°^C, 120 rpm, atmospheric air. For screening of NPQ induction and recovery phenotypes, cells were diluted to 100,000 cells mL^−1^ in a 24-well plate in fresh f/2 supplemented with 5 mM sodium bicarbonate (from a 1 M stock) to a final volume of 2 mL. F/2 was accompanied with 2 µl of either 100% ethanol (control) or 0.1, 0.5, 15, 10, 100 or 1000 nM final concentration β-estradiol (dissolved in 100% ethanol) serially diluted from a 10 mM stock. 24-well plates were placed in the same conditions. Cells were placed in 30 μmol photons m^−2^ s^−1^ white light conditions for 20 min prior to screening. An imaging PAM (MAXI head) was used to test for NPQ induction and relaxation properties. Fm and Fm’ were derived from an SP (840 ms, 3600 μmol photons m^−2^ s^−1^). The script used entailed an F_v_/F_m_ measurement, 5 min of 1250 μmol photons m^−2^ s^−1^ (SP every 30 s) and 15 min of 62 μmol photons m^−2^ s^−1^ (SP every 60 s for 3 min, subsequently every 120 s for 12 min).

### Statistics

Experiments were performed in biological replicates (n=3-6, see individual figure legends), except for Figure S2A. A repeated-measure one-way analysis of variance (RM-ANOVA) with a Tukey’s HSD was performed to test for significant variations between group means for most experiments, except for Figure 6, which used unpaired t-tests at each timepoint with Holm-Šídák’s multiple comparison method. Data represent mean values ± the standard deviation. Graphs and statistical analyses were performed in Prism version 10.1.2.

## Supplemental Data

**Supplemental Figure S1.** *Phaeodactylum* ZEP3 knockout design and genotype screening.

**Supplemental Figure S2.** NPQ and genotype screening for multiple ZEP3 complemented lines.

**Supplemental Figure S3.** Representative HPLC chromatograms from *Phaeodactylum* cultures during an NPQ induction experiment.

**Supplemental Figure S4.** Chlorophyll fluorescence and photo-physiological parameters of *Phaeodactylum* cultures during an NPQ induction experiment.

**Supplemental Figure S5.** Maximal specific growth rates of *Phaeodactylum* cultures during different constant light regimes.

**Supplemental Figure S6.** *Phaeodactylum* inducible ZEP3 strain design and genotype screening.

**Supplemental Figure S7.** Tuning of NPQ capacity via chemical induction of ZEP3 expression.

**Supplemental File 1** Gene sequences and Primers excel file.

## Author contributions

G.P., M.A.W. and Y.B. conceived the project and designed experiments. M.A.W., Y.B., A.J.P., T.K. and M.L. collected and analysed data. G.P. and M.A.W wrote the original draft of the manuscript with contributions from A.J.P. and M.L.. All authors contributed to review and editing of the manuscript and approved the final manuscript.

## Acknowledgements

We thank Michael Cantrell for early discussions about this work and Jesse Stahl for experimental assistance during strain screening.

## References

Alboresi A, Gerotto C, Giacometti GM, Bassi R, Morosinotto T (2010) *Physcomitrella patens* mutants affected on heat dissipation clarify the evolution of photoprotection mechanisms upon land colonization. Proc Natl Acad Sci USA 107: 11128–11133

Allahverdiyeva Y, Mustila H, Ermakova M, Bersanini L, Richaud P, Ajlani G, Battchikova N, Cournac L, Aro E-M (2013) Flavodiiron proteins Flv1 and Flv3 enable cyanobacterial growth and photosynthesis under fluctuating light. Proc Natl Acad Sci USA 110: 4111–4116

Allorent G, Lefebvre-Legendre L, Chappuis R, Kuntz M, Truong TB, Niyogi KK, Ulm R, Goldschmidt-Clermont M (2016) UV-B photoreceptor-mediated protection of the photosynthetic machinery in *Chlamydomonas reinhardtii*. Proc Natl Acad Sci USA 113: 14864–14869

Andersson B, Shen C, Cantrell M, Dandy DS, Peers G (2019) The Fluctuating Cell-Specific Light Environment and Its Effects on Cyanobacterial Physiology. Plant Physiology 181: 547–564

Arsalane W, Rousseau B, Duval J (1994) INFLUENCE OF THE POOL SIZE OF THE XANTHOPHYLL CYCLE ON THE EFFECTS OF LIGHT STRESS IN A DIATOM: COMPETITION BETWEEN PHOTOPROTECI’ION AND PHOTOINHIBITION. Photochem & Photobiology 60: 237–243

Bai Y, Cao T, Dautermann O, Buschbeck P, Cantrell MB, Chen Y, Lein CD, Shi X, Ware MA, Yang F, et al (2022) Green diatom mutants reveal an intricate biosynthetic pathway of fucoxanthin. Proc Natl Acad Sci USA 119: e2203708119

Bailleul B, Rogato A, De Martino A, Coesel S, Cardol P, Bowler C, Falciatore A, Finazzi G (2010) An atypical member of the light-harvesting complex stress-related protein family modulates diatom responses to light. Proc Natl Acad Sci USA 107: 18214–18219

Blommaert L, Chafai L, Bailleul B (2021) The fine-tuning of NPQ in diatoms relies on the regulation of both xanthophyll cycle enzymes. Sci Rep 11: 12750

Bonaventura C, Myers J (1969) Fluorescence and oxygen evolution from Chlorella pyrenoidosa. Biochimica et Biophysica Acta (BBA) - Bioenergetics 189: 366–383

Büchel C (2020) Light-Harvesting Complexes of Diatoms: Fucoxanthin-Chlorophyll Proteins. *In* AWD Larkum, AR Grossman, JA Raven, eds, Photosynthesis in Algae: Biochemical and Physiological Mechanisms. Springer International Publishing, Cham, pp 441–457

Buck JM, Sherman J, Bártulos CR, Serif M, Halder M, Henkel J, Falciatore A, Lavaud J, Gorbunov MY, Kroth PG, et al (2019) Lhcx proteins provide photoprotection via thermal dissipation of absorbed light in the diatom Phaeodactylum tricornutum. Nat Commun 10: 4167

Bugos RC, Yamamoto HY (1996) Molecular cloning of violaxanthin de-epoxidase from romaine lettuce and expression in Escherichia coli. Proc Natl Acad Sci USA 93: 6320–6325

Cao T, Bai Y, Buschbeck P, Tan Q, Cantrell MB, Chen Y, Jiang Y, Liu R-Z, Ries NK, Shi X, et al (2023) An unexpected hydratase synthesizes the green light-absorbing pigment fucoxanthin. The Plant Cell 35: 3053–3072

Cardol P, De Paepe R, Franck F, Forti G, Finazzi G (2010) The onset of NPQ and ΔμH+ upon illumination of tobacco plants studied through the influence of mitochondrial electron transport. Biochimica et Biophysica Acta (BBA) - Bioenergetics 1797: 177–188

Carter V, Rybicki NB (1990) Light Attenuation and Submersed Macrophyte Distribution in the Tidal Potomac River and Estuary. Estuaries 13: 441

Coesel S, Oborník M, Varela J, Falciatore A, Bowler C (2008) Evolutionary Origins and Functions of the Carotenoid Biosynthetic Pathway in Marine Diatoms. PLoS ONE 3: e2896

Dall’Osto L, Cazzaniga S, Wada M, Bassi R (2014) On the origin of a slowly reversible fluorescence decay component in the *Arabidopsis npq4* mutant. Phil Trans R Soc B 369: 20130221

Dautermann O, Lyska D, Andersen-Ranberg J, Becker M, Fröhlich-Nowoisky J, Gartmann H, Krämer LC, Mayr K, Pieper D, Rij LM, et al (2020) An algal enzyme required for biosynthesis of the most abundant marine carotenoids. Sci Adv 6: eaaw9183

De Souza AP, Burgess SJ, Doran L, Hansen J, Manukyan L, Maryn N, Gotarkar D, Leonelli L, Niyogi KK, Long SP (2022) Soybean photosynthesis and crop yield are improved by accelerating recovery from photoprotection. Science 377: 851–854

Gerotto C, Morosinotto T (2013) Evolution of photoprotection mechanisms upon land colonization: evidence of PSBS -dependent NPQ in late Streptophyte algae. Physiologia Plantarum 149: 583–598

Giovagnetti V, Flori S, Tramontano F, Lavaud J, Brunet C (2014) The Velocity of Light Intensity Increase Modulates the Photoprotective Response in Coastal Diatoms. PLoS ONE 9: e103782

Giovagnetti V, Ruban AV (2017) Detachment of the fucoxanthin chlorophyll a / c binding protein (FCP) antenna is not involved in the acclimative regulation of photoprotection in the pennate diatom Phaeodactylum tricornutum. Biochimica et Biophysica Acta (BBA) - Bioenergetics 1858: 218–230

Gorbunov MY, Falkowski PG (2022) Using Chlorophyll Fluorescence to Determine the Fate of Photons Absorbed by Phytoplankton in the World’s Oceans. Annu Rev Mar Sci 14: 213–238

Jahns P, Latowski D, Strzalka K (2009) Mechanism and regulation of the violaxanthin cycle: The role of antenna proteins and membrane lipids. Biochimica et Biophysica Acta (BBA) - Bioenergetics 1787: 3–14

Jallet D, Caballero MA, Gallina AA, Youngblood M, Peers G (2016) Photosynthetic physiology and biomass partitioning in the model diatom Phaeodactylum tricornutum grown in a sinusoidal light regime. Algal Research-Biomass Biofuels and Bioproducts 18: 51–60

Jiang Y, Cao T, Yang Y, Zhang H, Zhang J, Li X (2023) A chlorophyll *c* synthase widely co-opted by phytoplankton. Science 382: 92–98

Jinkerson RE, Poveda-Huertes D, Cooney EC, Cho A, Ochoa-Fernandez R, Keeling PJ, Xiang T, Andersen-Ranberg J (2024) Biosynthesis of chlorophyll c in a dinoflagellate and heterologous production in planta. Current Biology 34: 594–605.e4

Kassaw TK, Paton AJ, Peers G (2022) Episome-Based Gene Expression Modulation Platform in the Model Diatom Phaeodactylum tricornutum. ACS Synth Biol 11: 191–204

Krause GH (1988) Photoinhibition of photosynthesis. An evaluation of damaging and protective mechanisms. Physiologia Plantarum 74: 566–574

Kromdijk J, Glowacka K, Leonelli L, Gabilly S, Iwai M, Niyogi K, Long S (2016) Improving photosynthesis and crop productivity by accelerating recovery from photoprotection. SCIENCE 354: 857–861

Külheim C, Ågren J, Jansson S (2002) Rapid Regulation of Light Harvesting and Plant Fitness in the Field. Science 297: 91–93

Kunugi M, Satoh S, Ihara K, Shibata K, Yamagishi Y, Kogame K, Obokata J, Takabayashi A, Tanaka A (2016) Evolution of Green Plants Accompanied Changes in Light-Harvesting Systems. Plant Cell Physiol 57: 1231–1243

Kunz H-H, Gierth M, Herdean A, Satoh-Cruz M, Kramer DM, Spetea C, Schroeder JI (2014) Plastidial transporters KEA1, -2, and -3 are essential for chloroplast osmoregulation, integrity, and pH regulation in *Arabidopsis*. Proc Natl Acad Sci USA 111: 7480–7485

Lavaud J, Materna AC, Sturm S, Vugrinec S, Kroth PG (2012) Silencing of the Violaxanthin De-Epoxidase Gene in the Diatom Phaeodactylum tricornutum Reduces Diatoxanthin Synthesis and Non-Photochemical Quenching. PLoS ONE 7: e36806

Lavaud J, Rousseau B, Etienne A-L (2002) In diatoms, a transthylakoid proton gradient alone is not sufficient to induce a non-photochemical fluorescence quenching. FEBS Letters 523: 163–166

Lohr M, Wilhelm C (1999) Algae displaying the diadinoxanthin cycle also possess the violaxanthin cycle. Proc Natl Acad Sci USA 96: 8784–8789

MacIntyre HL, Kana TM, Anning T, Geider RJ (2002) Photoacclimation of photosynthesis irradiance response curves and photosynthetic pigments in microalgae and cyanobacteria. Journal of Phycology 38: 17–38

Malnoë A, Schultink A, Shahrasbi S, Rumeau D, Havaux M, Niyogi KK (2018) The Plastid Lipocalin LCNP Is Required for Sustained Photoprotective Energy Dissipation in Arabidopsis. Plant Cell 30: 196–208

Marin E, Nussaume L, Quesada A, Gonneau M, Sotta B, Hugueney P, Frey A, Marion-Poll A (1996) Molecular identification of zeaxanthin epoxidase of Nicotiana plumbaginifolia, a gene involved in abscisic acid biosynthesis and corresponding to the ABA locus of Arabidopsis thaliana. EMBO J 15: 2331–2342

Maxwell K, Johnson GN (2000) Chlorophyll fluorescence - a practical guide. Journal of Experimental Botany 51: 659–668

Murchie EH, Ruban AV (2020) Dynamic non-photochemical quenching in plants: from molecular mechanism to productivity. The Plant Journal 101: 885–896

Nawrocki WJ, Buchert F, Joliot P, Rappaport F, Bailleul B, Wollman F-A (2019) Chlororespiration Controls Growth Under Intermittent Light. Plant Physiol 179: 630–639

Nilkens M, Kress E, Lambrev P, Miloslavina Y, Müller M, Holzwarth AR, Jahns P (2010) Identification of a slowly inducible zeaxanthin-dependent component of non-photochemical quenching of chlorophyll fluorescence generated under steady-state conditions in Arabidopsis. Biochimica et Biophysica Acta (BBA) - Bioenergetics 1797: 466–475

Nymark M, Valle KC, Hancke K, Winge P, Andresen K, Johnsen G, Bones AM, Brembu T (2013) Molecular and Photosynthetic Responses to Prolonged Darkness and Subsequent Acclimation to Re-Illumination in the Diatom Phaeodactylum tricornutum. PLoS ONE 8: e58722

Peers G, Truong TB, Ostendorf E, Busch A, Elrad D, Grossman AR, Hippler M, Niyogi KK (2009) An ancient light-harvesting protein is critical for the regulation of algal photosynthesis. Nature 462: 518–U215

Peers, G (2015) Enhancement of biomass production by disruption of light energy dissipation pathways. U.S. Patent No. 8,940,508. Washington, DC: Patent and Trademark Office.

Perin G, Bellan A, Michelberger T, Lyska D, Wakao S, Niyogi KK, Morosinotto T (2023) Modulation of xanthophyll cycle impacts biomass productivity in the marine microalga *Nannochloropsis*. Proc Natl Acad Sci USA 120: e2214119120

Ritchie RJ (2008) Universal chlorophyll equations for estimating chlorophylls a, b, c, and d and total chlorophylls in natural assemblages of photosynthetic organisms using acetone, methanol, or ethanol solvents. Photosynthetica 46: 115–126

Ruban A, Lavaud J, Rousseau B, Guglielmi G, Horton P, Etienne A-L (2004) The super-excess energy dissipation in diatom algae: comparative analysis with higher plants. Photosynth Res 82: 165–175

Ruban AV, Berera R, Ilioaia C, Van Stokkum IHM, Kennis JTM, Pascal AA, Van Amerongen H, Robert B, Horton P, Van Grondelle R (2007) Identification of a mechanism of photoprotective energy dissipation in higher plants. Nature 450: 575–578

Sacharz J, Giovagnetti V, Ungerer P, Mastroianni G, Ruban AV (2017) The xanthophyll cycle affects reversible interactions between PsbS and light-harvesting complex II to control non-photochemical quenching. Nature Plants 3: 16225

Takaichi S (2011) Carotenoids in Algae: Distributions, Biosyntheses and Functions. Marine Drugs 9: 1101–1118

Ware MA, Dall’Osto L, Ruban AV (2016) An In Vivo Quantitative Comparison of Photoprotection in Arabidopsis Xanthophyll Mutants. Front Plant Sci. doi: 10.3389/fpls.2016.00841

Ware MA, Hunstiger D, Cantrell M, Peers G (2020) A Chlorophyte Alga Utilizes Alternative Electron Transport for Primary Photoprotection. Plant Physiology 183: 1735–1748

Wraight CA, Crofts AR (1970) Energy-Dependent Quenching of Chlorophyll *a* Fluorescence in Isolated Chloroplasts. European Journal of Biochemistry 17: 319–327

